# Bioengineering a novel 3D in-vitro model to recreate physiological oxygen levels and tumor-immune interactions

**DOI:** 10.1101/828145

**Authors:** Somshuvra Bhattacharya, Kristin Calar, Claire Evans, Mark Petrasko, Pilar de la Puente

## Abstract

Oxygen deprivation within tumors is one of the most prevalent causes of resilient cancer cell survival and increased immune evasion in breast cancer (BCa). Current *in vitro* models do not adequately mimic physiological oxygen levels relevant to breast tissue and its tumor-immune interactions. Here, we propose an approach to engineer a three-dimensional (3D) model (named 3D engineered oxygen, 3D-O) that supports growth of BCa cells and generates physio- and pathophysiological oxygen levels. Low oxygen-induced changes within the 3D-O model supported known tumor hypoxia characteristics such as reduced BCa cell proliferation, increased extracellular matrix protein expression, increased extracellular vesicle secretion and enhanced immune surface marker expression on BCa cells. We further demonstrated that low oxygen-induced changes mimicked tumor-immune interactions leading to immune evasion mechanisms. CD8+ T cell infiltration was significantly impaired under pathophysiological oxygen levels and we were able to establish that hypoxia inhibition re-sensitize BCa cells to cytotoxic CD8+ T cells. Therefore, our novel 3D-O model could serve as a promising platform for the evaluation of immunological events and as a drug-screening platform tool to overcome hypoxia-driven immune evasion.

## INTRODUCTION

With 1 million new cases in the world every year, breast cancer (BCa), excluding skin cancer, is the most common cancer in women.(1, 2). Close to 1 in 8 women will be diagnosed with BCa in their lifetime, accounting for almost 18% of all cancer in women (3). As of 2018, the American Cancer Society estimates that each year about 2,000 new cases of invasive BCa are being diagnosed (4). Despite breast tumors being more manageable now, mortality from BCa resistance, recurrence and metastasis is still the number one challenge that advancements in BCa therapy have consistently failed to eliminate (5). Cancer immune surveillance consists of the ability of the immune system to recognize and eliminate tumor cells (6). BCa tumor cells avoid tumor corrective processes intrinsically executed by the patient’s host immunity hence undergoing sustained survival (7). While a demonstrated durable response to immunotherapeutic intervention has been shown in some types of cancer, such as melanoma, bladder, and renal cell carcinoma, BCa has not shown the same efficacy (8).

In regards to this, many studies have been directed towards understanding the mechanisms of immune evasion in BCa (9–11). The knowledge gained from some of these studies has collectively generated many improved immune-therapeutic leads which function either by boosting a patient’s intrinsic immune arsenal e.g. immune checkpoint inhibitions (12) or recovering immune operational performance by reducing the influence of immune-suppressive molecules generated by the BCa tumor (13). Increasing evidence suggests that regulation of immune evasion in BCa extends well beyond the authority of merely resident tumor cells. BCa tumor microenvironment (TME) parameters such as extracellular matrix (ECM), cell-matrix interactions, growth factors, cytokines, and an oxygen-deprived niche can heavily contribute to immune evasion as well (14–17).

Intratumoral hypoxia, defined by the deprivation of oxygen within tumors, has been associated with modified cellular expression levels, altered ECM secretion, and active immune evasion (18–21). Patients with tumors that are poorly oxygenated have an increased risk of mortality (22). In addition, the presence of tumor-infiltrated lymphocytes (TILs) can be used as a relevant prognostic indicator (23–25). A notorious hallmark associated with BCa intratumoral oxygen deprivation is alteration in the cancer cell survival metabolism to function through the hypoxia-inducible factor-1α (HIF-1α) regulatory pathway (26–28). Physiological oxygen content can vary widely from tissue to tissue, and with healthy breast having been reported to harbor a partial pressure oxygen (pO_2_) content range from approximately 55 mm Hg i.e. 7.33 kPa to 85 mm Hg or 11 kPa (29). Oxygen deprivation within the BCa tumor can take this content down to almost 1.3 kPa, and approximately 30% of BCa tumors even exhibit pO_2_ values less than 0.266 kPa (29–31).

In the efforts to recreate oxygen deprivation-driven tumor immune evasion mechanisms in the laboratory, very few cancer *in-vitro* models adequately mimic physiological oxygen levels relevant to breast tissue and its tumor-immune interactions (32–34). Traditional two-dimensional (2D) culture models fail to generate oxygen gradients as such, and hence experiments using these models expose the cells to higher than physiological oxygen levels (35). These models might not accurately demonstrate tumor-immune evasive outcomes. In order to overcome these limitations, relevant three-dimensional (3D) culture models have been studied. A wide array of matrices, including synthetic and natural, have been developed to recapitulate key features of the TME (36–41). While biochemical and physical parameters, such as conduciveness to key biochemical signals, stiffness, degradability, permeability to nutrients, diffusibility to gases and swelling indices have been heavily studied (42–48), recreation of physiological oxygen levels and their influence on tumor-immune interactions remains fairly under investigated.

Therefore, the purpose of our study is to create a novel tissue culture model that supports growth of BCa cells and generates physio- and pathophysiological oxygen levels. In this regard, we bioengineered a novel 3D *in-vitro* model to recreate physio- and pathophysiological oxygen levels and tumor-immune interactions, named 3D engineered oxygen (3D-O) model. A proprietary 3D human blood plasma matrix is used as the framework upon which the 3D culture exists. We hypothesize that the 3D-O model will appropriately mimic relevant oxygen conditions of the breast microenvironment, while serving as a highly maneuverable platform that can simulate the intrinsic biological changes associated with the oxygen-deprived tumor environment where the breast cancer cells reside, and further investigate the prevailing low oxygen-driven consequences in tumor-immune interactions.

## MATERIALS AND METHODS

### Reagents

Calcium chloride (CaCl_2_), trans-4-(Aminomethyl)cyclohexanecarboxylicacid (AMCHA),, dimethyl sulfoxide (DMSO), Ficoll-Paque density gradient medium, DAPI, and glutaraldehyde, were purchased from Sigma-Aldrich (Saint Louis, MO). Type I collagenase and Image-iT™ Green Hypoxia detection reagent and Triton X-100 were purchased from Thermo Fischer Scientific, (Waltham, MA). Cell tracker DiO (excitation, 488 nm; emission, 525/50 nm) was purchased from Invitrogen (Carlsbad, CA). Drugs including PX-478 and Durvalumab were purchased from Selleck Chemicals (Houston, TX).

### Cell lines

The BCa cell lines (MDA-MB-231 and MCF-7) were kind gifts from Dr. Kristi Egland (Sanford Research, Sioux Falls, SD). All human cell lines were authenticated by short tandem repeat profiling (Genetica DNA Laboratories, Cincinnati, OH). In addition, all cell lines were confirmed mycoplasma free. All cell lines were cultured at 37°C, 5% CO_2_ in DMEM media (Corning CellGro, Mediatech, Manassas, VA) which was supplemented with 10% fetal bovine serum (FBS, GiBCo, Life technologies, Grand island, NY), 100 U/ml penicillin, and 100 μg/ml streptomycin (Corning CellGro, VA). Before experiments, in some cases, BCa cells (1 × 10^6^ cells/ml) were prelabeled with DiO (10 μg/ml) for 1 hour.

### Primary cells

Primary peripheral blood mononuclear cells (PBMCs) were isolated from healthy blood provided by the Sanford USD Medical Center, Sioux Falls, SD using SepMate PBMC isolation tubes (STEMCELL Technologies, Canada) following manufacturer instructions. Informed consent was obtained from all healthy subjects with an approval from the Sanford Health Institutional Review Board and in accordance with the Declaration of Helsinki. Collected fresh blood was first centrifuged at 3000 rpm for 10 minutes at room temperature, to separate plasma. After plasma isolation, leftover blood was then diluted with phosphate-buffered saline (PBS) containing 2% FBS in equal volume to blood and pipetted slowly along the wall of the SepMate tube in which Ficoll-Paque density gradient medium was added beforehand. After 10 minutes of centrifugation with the brake on, highly purified PBMCs were poured into a new tube. Finally, the cells were washed three times by centrifuging at 100 × g for 10 minutes at room temperature and suspending in PBS containing 2% FBS solution for each wash cycle. The enriched cells were counted and cells were viably frozen at −80° in FBS with 10% (v/v) DMSO. PBMCs were pre-analyzed by flow cytometry to determine the frequency of immune cell population before using them in any experiment to account for variability between patients.

### Development of 3D-O cultures

Using proprietary bioengineering methods, the 3D-O matrix can be engineered through the cross-linking of fibrinogen into fibrin. Calcium chloride (CaCl_2_) and AMCHA were used as a cross-linker and stabilizer, respectively. A mixture of plasma, BCa cell suspension (30×10^4^ cells/per scaffold) in DMEM complete media, AMCHA and CaCl_2_ was prepared with a 4:4:1:1 volume ratio, respectively. In order to avoid intra-assay variability due to the matrix, plasma was pooled in batches. Matrices were prepared in 96 well plates or in 8-well chamber slides for confocal imaging. Matrices were allowed to stabilize in an incubator at 37°C, before being covered with additional DMEM complete media. After gelification was completed, 3D-O matrices were incubated for 4 days at 37°C, while being exposed to variable O_2_ environments (21%, 5%, and 1.5% O_2_). In the case of tumor-immune infiltration assays, at day 4, PBMCs were incorporated as a cell suspension in the medium added on the top of the matrix (1:3 to 1:6 cancer: PBMCs ratio) and kept at 37°C, while being exposed to the same O_2_ environment up to day 7 (**Figure 1A**).

**Figure 1:**
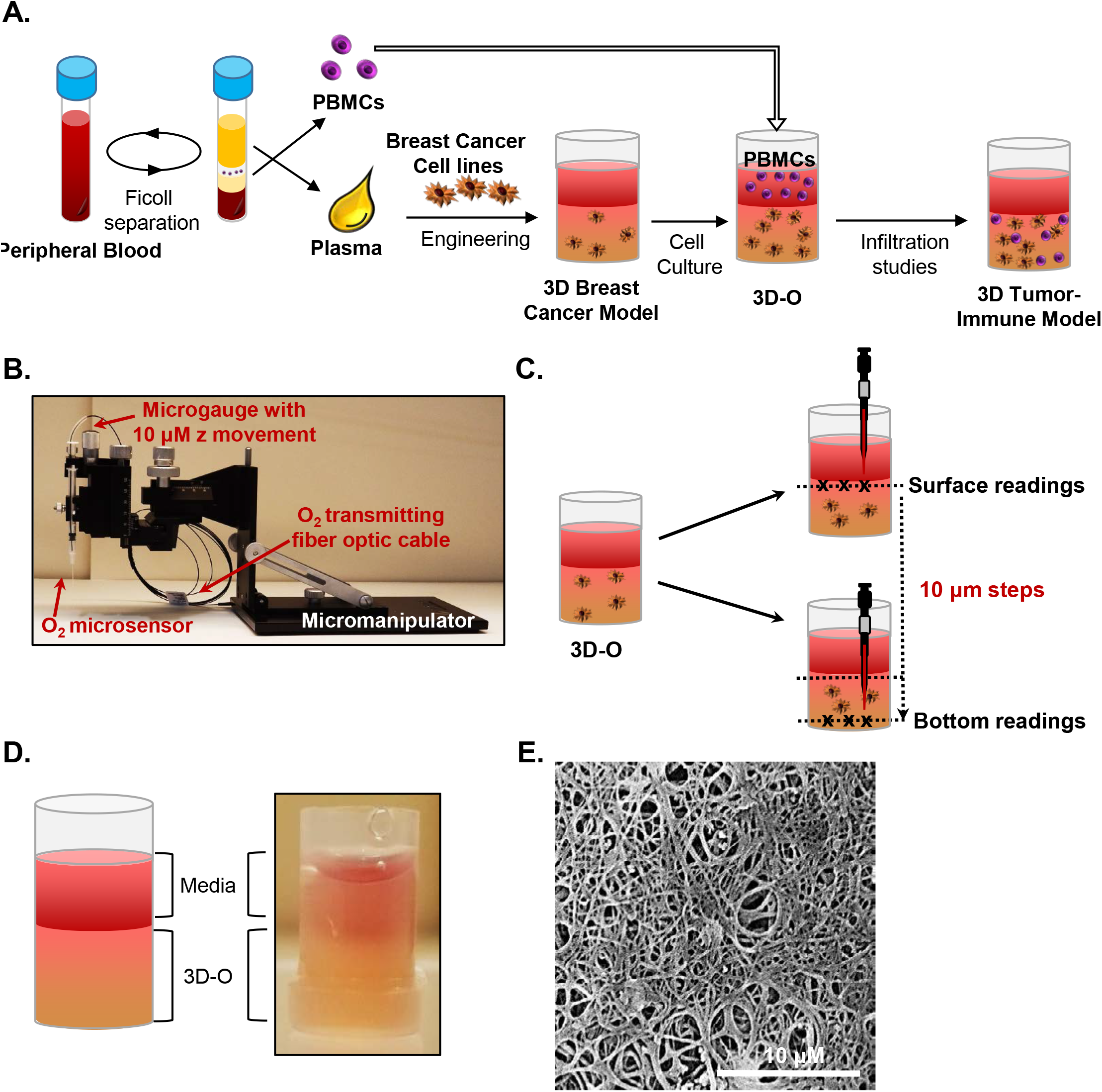
Methodology of 3D-O matrices. A) 3D-O matrices were developed through cross-linking of fibrinogen (naturally found in the plasma of blood supernatant) including BCa cells. PBMCs were added on top of the 3D-O scaffolds on day 4 of culture and allowed to infiltrate into the scaffold until day 7. B) Oxygen microsensor (PreSens) and Manual Micromanipulator configuration for O_2_ profiling in the Z-direction every 10μm. C) Oxygen microsensor was extended delicately and safely with 10 μm profiling accuracy to determine three surface (border between media and 3D-O matrix) and bottom (bottom of the well) readings. D) Visual and schematic representation of a 3D-O matrix with medium on top. E) SEM micrograph of acellular 3D-O scaffold cultured on day 4. Scale bar: 10 μm.

### Oxygen measurement within 3D-O matrices

Oxygen partial pressure (pO_2_) levels were measured in acellular and cell-seeded 3D-O matrices. 3D-O scaffolds were profiled along the z-direction with an oxygen microsensor (Needle-Type Oxygen Microsensor NTH-PSt7, PreSens, Regensburg, Germany) and a manual micromanipulator (**Figure 1B**). Oxygen measurement by this type of sensor has been described previously (49–51). Briefly, to record oxygen pressure, the sensor was introduced into the geometric center (3 measure points) of the 3D-O and moved from the border between the media and 3D-O (Surface) in 10μm steps towards the bottom of the well plate (Bottom), as illustrated in **Figure 1C**. The Software, PreSens Profiling Studio, enabled the measurement of variable step sizes, measuring velocities and wait times. Before application, a two-point calibration was performed: 100% CO_2_ as 0% O_2_ reference and ambient air as 21% O_2_ reference. Increasing matrix height, incubation time, and variable O_2_ incubation environments (21% O_2_ to 1.5% O_2_) were investigated.

### Scanning electron microscopy (SEM) studies

Samples for SEM analysis were subjected to gradual cooling (−20°C to −80°C) after fixation with 10% glutaraldehyde solution for 6-8 hours. After the fixed samples were completely frozen, the samples were subjected to lyophilization for 48 hours at about −105°C/100 mTorr vacuum. Freeze-dried samples were attached to an aluminum sample stub with conductive silver paint (Ted Pella, Inc., Redding, CA) and tightened using a copper tape. Samples were then subjected to sputter-coating with a layer of ~10 nm gold film before imaging by SEM. SEM images were acquired using a FEI Quanta 450 Scanning Electron Microscope at different magnifications. The accelerating voltage used was 10 kV, and images were acquired using a dwell time of 30-60 μs.

### Immunohistochemistry (IHC) and Immunofluorescence (IF) studies

Acellular or 3D-O matrices containing BCa cells (cell-seeded) were fixed in 10% neutral buffered formalin and processed on a Leica 300 ASP tissue processor. The matrices were oriented on the slide so that the top of the matrix section always faced forward. Paraffin embedded matrix sections were longitudinally sliced at 10 μm. The BenchMark® XT automated slide staining system (Ventana Medical Systems, Inc., AZ) was used for the antibody optimization and staining. The antigen retrieval step was performed using the Ventana CC1 solution, which is a basic pH tris based buffer. Both primary and secondary antibodies were prepared in a 1X permeabilization buffer (BioLegend, CA). The Ventana iView DAB detection kit was used as the chromogen and the slides were counterstained with hematoxylin. Anti-Ki-67 (CRM325, 1:100, Biocare Medical) and anti-CD8 (MA5-14548, 1:100, Invitrogen, CA) primary antibodies were used. IHC images were imaged using an Aperio VERSA Bright field Fluorescence & FISH Digital Pathology Scanner (Leica, NJ). Omission of the primary antibody served as the negative control. Secondary antibodies used were biotin-conjugated goat anti-Rabbit IgG (111-065-144, 1:1000, Jackson ImmunoResearch, PA) and biotin-conjugated donkey anti-mouse IgG (715-065-151, 1:1000, Jackson ImmunoResearch, PA), respectively. For IF studies, paraffin sections were dewaxed in the following order: 10 minutes in xylene, 10 minutes in 100% ethanol, 10 minutes in 95% ethanol, 10 minutes in 70% ethanol and 10 minutes in distilled water, followed by rehydration in wash buffer (0.02% BSA in PBS) for 10 minutes. After this, sections were subjected to incubation in blocking buffer (5% BSA in PBS) for 60 minutes at room temperature to block non-specific staining between the primary antibodies and the sample. Sections were rinsed with washing buffer and incubated in incubation buffer (1% BSA in PBS) with different primary antibodies. Primary antibody incubation was carried out overnight at 4°C to allow for optimal binding of antibodies to sample targets and reduce non-specific background staining. Anti-collagen-I (MA1-26771, 1:100, Thermo Fischer Scientific, MA), anti-collagen-III (SAB4200749, 1:100, Sigma Aldrich, MO), anti-laminin (SAB4200719, 1:100, Sigma Aldrich, MO), anti-fibronectin (SAB4200784, 1:100, Sigma Aldrich, MO) and an AlexaFluor 647 conjugated anti-HIF-1α were used (359706, 1:100, Biolegend, CA). A FITC conjugated secondary antibody (SAB4600042, 1:1000, Sigma Aldrich, MO) was used whenever applicable. For samples stained with anti-HIF-1α, blocking and incubation buffers were prepared in 1X permeabilization buffer (Biolegend, CA). Dilution of antibodies were carried out according to manufacturer’s instructions. Lastly, a drop of anti-fade mounting media containing DAPI was added to the samples and sections were imaged.

### Confocal imaging

3D-O matrices containing BCa cells alone or in co-culture with PBMCs growing in 8 well chambers and paraffin section cuts of 3D-O matrices were imaged using a Nikon Ti2-A1TR confocal microscope (×20 dry, ×40 oil and ×60 oil objectives, 2.5 magnified) and analyzed using NIS elements software (Nikon, Melville, NY, USA). Oxygen deprivation was imaged by using Image-iT™ Green Hypoxia reagent (5μM) on the day of analysis. Samples were excited at 488 nm (FITC/DiO/ Image-iT™ Green Hypoxia Reagent), 358 nm (DAPI), 640 nm (APC-Cy7) and the emission light was collected at 500 – 530 nm, 461 nm, and 650 nm long pass, for each channel respectively. Z-stack images of approximately 0.8 mm thickness were taken for each sample at 2 μm step sizes. Each frame consisted of a 520 × 520 pixel image, taken at a rate of 1 μs/pixel.

### Extracellular vesicle isolation and characterization

Extracellular vesicle isolation and their characterization was performed as previously described (52–54) using similar protocols. Briefly, 3D-Os were seeded with MDA-MB-231 BCa cells and cultured for 4 days in DMEM medium containing 10% FBS that was depleted of exosomes. FBS was made extracellular vesicle free by performing an overnight ultracentrifugation at 100,000 × g. Conditioned medium samples were collected after 4 days and extracellular vesicles were purified by differential ultracentrifugation: first, samples were centrifuged at 300 × g for 10 minutes at 4°C to pellet out the cells. Then, the supernatant was centrifuged at 2,000 × g for 20 minutes at 4°C, followed by transferring to a new tube, and then centrifugation for 30 minutes at 10,000 × g. Finally, the samples were subjected to a final spin using a SureSpin 630/17 rotor for 120 minutes at 100,000 × g. All pellets were washed in PBS and re-centrifuged at the same speed and re-suspended in 400 μL of sterile PBS. After this, the BCA protein assay kit (Thermo Fischer Scientific, MA) was utilized with some modifications to compensate for the low protein yield from extracellular vesicle preparations. Briefly, 5 μl of 10% Triton X-100 was added to an aliquot of 50 μl of purified 3D-O extracellular vesicles and incubated 10 minutes at room temperature. A 1:11 ratio of sample to working reagent was used and incubated in a 96-well plate for 1 hour at 37°C. Absorbance at 562 nm was then assessed using a SpectraMax Plus 384 (Molecular Devices, CA) and protein concentration estimated from a quartic model fit to the BSA standard curve. Finally, the purified vesicles were characterized using the NanoSight particle analyzer (Nanosight, UK), following the manufacturer’s protocol.

### BCa proliferation, hypoxic status and surface marker expression using flow cytometry

3D-O matrices were enzymatically digested with collagenase (20 mg/ml for 2 - 3 hours at 37°C) on day 4. BCa cells were isolated and identified by gating cells with a high DiO signal (excitation, 488 nm; emission, 530/30 nm). Antibodies used to evaluate hypoxic status and surface marker expression were AlexaFluor 647 conjugated anti-HIF-1α (359706, Biolegend, CA), APC-Cy7 conjugated anti-CD8 (344714, Biolegend, CA), PE conjugated anti-PD-L1 (393608, Biolegend, CA), PE conjugated anti-MUC-1 (355608, Biolegend, CA), and PerCP-Cy5 conjugated anti-CD73 (344014, Biolegend, CA). Cell viability was evaluated by using a Sytox Blue live-dead fluorescent dye (S34857, Invitrogen, CA) possessing excitation, 358 nm; emission, 461 nm or Live/Dead Blue cell stain (L34962, Thermo Fischer Scientific, MA). For all analysis, a minimum of 5,000 events were acquired using BD FACS Fortessa and FACSDiva v6.1.2 software or BD FACS Accuri and BS Accuri C6 software (BD Biosciences), respectively. The BCa cell counts were always normalized to a predetermined number of counting beads (424902, Biolegend, CA), and mean fluorescence intensity (MFI) ratios for each of the above mentioned targets studied were assessed with respect to the corresponding isotype in the BCa-DiO^+^ cells. The data was analyzed using FlowJo program v10 (Ashland, OR).

### Lymphocyte infiltration characterization

Differences in lymphocyte infiltration into 3D-O scaffolds as a result of different oxygen gradients were assessed. PBMCs were incorporated as cell suspension in the medium added on the top of the matrix at day 4 and analyzed at day 7. 3D-O matrices were enzymatically digested with collagenase and PBMCs were isolated and surface-stained with the following antibodies: FITC conjugated anti-CD3 (300406, Biolegend, CA), PE-Cy5 conjugated anti-CD4 (300508, Biolegend, CA), APC-Cy7 conjugated anti-CD8 (344714, Biolegend, CA), APC conjugated anti-CD19 (302212, Biolegend, CA) and BV510 conjugated anti-CD45 (304036, Biolegend, CA). Infiltrated populations were characterized with manual gating, or combined datasets were down-sampled and subjected to dimensionality reduction using t-stochastic neighbor embedding (t-SNE) algorithm (55, 56) or automatically defined with FlowSOM clustering algorithm (57). Mean absolute numbers of CD3+, CD4+, CD8+ and CD19+ cells (normalized to beads) were determined in each experimental group. To confirm differences in CD8+ infiltration as a result of oxygen gradients, number of infiltrated CD8+ cells were imaged using confocal microscopy or IHC.

### Effect of reversing oxygen deprivation within 3D-O matrices

To attribute oxygen deprivation as the primary intermediary for all observed differences, we used a HIF inhibitor as a single agent (PX-478, 5 μM). PX-478 inhibitor was tested in HIF-1α expression, BCa cell proliferation, collagen-I expression, and CD8+ infiltration. We further tested a selective, high affinity human IgG1 mAb that blocks programmed cell death ligand-1 (PD-L1) binding to PD-1 (Durvalumab, 5 μM) to evaluate its role on CD8 infiltration by flow cytometry as previously described.

### Statistical Analysis

Experiments were performed in triplicates and repeated at least three times. Results are presented as mean ± standard deviation, and statistical significance was analyzed using student t-test or one-way ANOVA; a p-value less than 0.05 was considered significant. Outliers were removed using Interquartile Range (IQR).

## RESULTS

### 3D-O matrices demonstrate development of oxygen gradients

Plasma was engineered to create a gelatinous-like (**Figure 1D**) and porous (**Figure 1E**) 3D-O matrix. pO_2_ levels were profiled from surface to bottom inside 3D-O models to evaluate the effects of height, time of incubation and low oxygen induced by incubation in a hypoxic incubator in cell-seeded and acellular 3D-O models. 3D-O height, incubation time, presence of BCa cells and surrounding oxygen content were observed to influence oxygen levels within its niche. As illustrated in **Figure 2Ai**, acellular 3D-O scaffolds, after incubation in 21% O_2_ for increased incubation time, did not exhibit significant development of pO_2_ gradients with the lowest values recorded at bottom 19.39 kPa, 18.25 kPa and 16.97 kPa for days 2, 4 and 7, respectively. In contrast, 3D-O matrices were much more dynamic in generating pO_2_ gradients when exposed to 1.5% O_2_ for increasing incubation times, especially for the 3D-O acellular scaffold incubated for 7 days, which showed a steep drop from 19.76 kPa at the surface to 8.49 kPa at the bottom (**Figure 2Aii**). When matrix height was studied, we found that while 3D-O matrices incubated at 21% O_2_ for 4 days did not substantially change pO_2_ levels in either the surface or bottom (**Figure 2Bi**), 3D-O matrices incubated at 1.5% O_2_ showed a gradual drop in pO_2_ levels from surface to bottom with increasing heights (**Figure 2Bii**). While 2 mm height 3D-O matrices generate gradients from 18.7kPa to 15.6kPa, increasing heights decreased the oxygen levels to 12.7kPa and 8.3 kPa for surface and bottom in 4 mm matrices, respectively, confirming that oxygen levels can be controlled and manipulated in 3D-O matrices.

**Figure 2:**
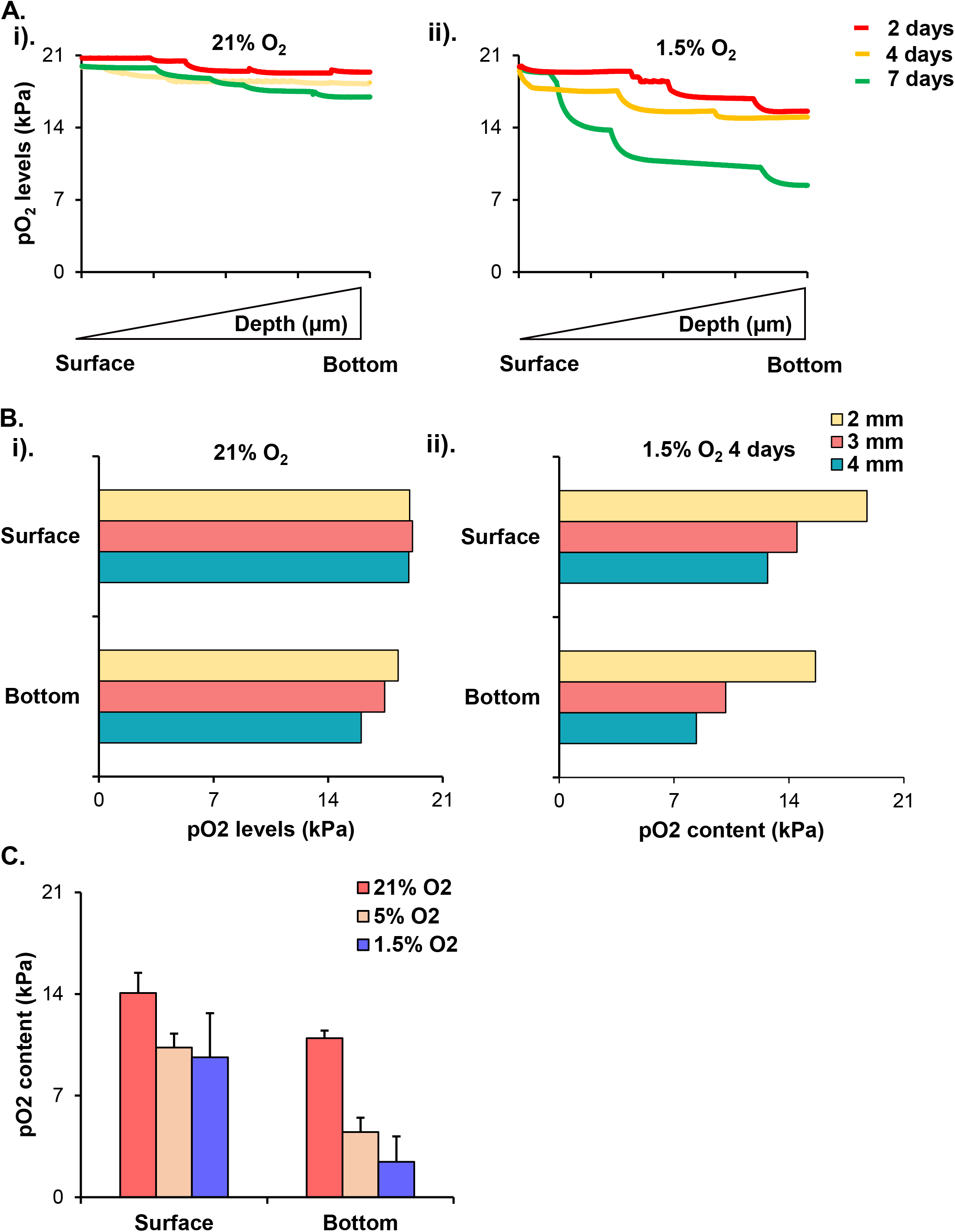
3D-O matrices demonstrate the generation of physiologically relevant oxygen levels. A) Surface to bottom pO_2_ levels (kPa) for acellular 3D-O matrices incubated for 2, 4, and 7 days either under 21% O_2_ (i) or 1.5% O_2_ (ii). B) Surface to bottom pO_2_ levels (kPa) for acellular 3D-O matrices of different heights (2, 3, 4 mm) incubated for 4 days either under 21% O_2_ (i) or 1.5% O_2_ (ii). C) Surface to bottom pO_2_ levels (kPa) for cell-seeded 3D-O matrices incubated for 4 days either under 21%, 5% and 1.5% O_2_.

Furthermore, the cell-seeded 3D-O matrices demonstrated wider pO_2_ gradients compared to the acellular samples at the same height, incubation time and oxygen incubation. Cell-seeded 3D-O matrices incubated at 21% O_2_ were found to exhibit a surface pO_2_ value of 14.06 kPa and a bottom value of 10.95 kPa (**Figure 2C**). A gradual decrease in surface and bottom pO_2_ levels was manipulated by oxygen incubation in hypoxic chamber. When incubated at 5% O_2_, 3D-O matrices containing BCa cells exhibited pO_2_ values of 10.3 at the surface to 4.5 kPa at the bottom. Simultaneously, when the scaffolds were incubated at 1.5% O_2_ the pO_2_ levels developed within the scaffolds dropped to 9.64 kPa and2.4 kPa at surface and bottom, respectively (**Figure 2C**). Hereafter, the 3D-O matrices will be referred to as 3D-O physiological (gradient 14 – 10 kPa) and 3D-O tumorous (gradient 9.6 to 2.4 kPa), respectively.

The existence of the oxygen gradient across the depth of 3D-O containing BCa cells was confirmed using confocal microscopy and the hypoxia-sensing dye (Image-iT Hypoxia Reagent). 3D-O tumorous matrices showed a significant increase in the number of BCa cells expressing HIF-1α, in which the MFI ratio was 1.6 and 3.7 times higher compared to 3D-O physiological matrices for MDA-MD-231 and MCF7, respectively (**Figure 3Ai**). Representative flow cytometry histograms of Alexafluor 647-HIF-1α signal in 3D-O physiological and 3D-O tumorous compared to isotype control BCa cells showed an increase HIF-1α expression in 3D-O tumorous compared to 3D-O physiological matrices (**Figure 3Aii**). Quantification of HIF-1α signal in bottom areas of 3D-O physiological and 3D-O tumorous revealed significant differences. While the top areas of the cultures showed few BCa cells expressing HIF-1α for both conditions, deeper areas in the bottom of 3-O tumorous matrices showed an increased number of BCa cells expressing HIF-1α compared to 3D-O physiological matrices (**Figure 3B and C**). A hypoxia reagent that fluoresces when oxygen levels fall below 5%, confirmed that greater number of BCa cells growing within 3D-O tumorous matrices exhibited positive green fluorescent signal compared to the 3D-O physiological matrices, mainly at the bottom areas (**Figure 3D**). We then evaluated the effect of HIF inhibitor PX-478 (5 μM) on HIF-1α expression under 3D-O physiological or tumorous conditions. PX-478 decreased the expression of HIF-1α in BCa cells incubated in 3D-O tumorous conditions to the level of 3D-O physiological conditions (**Figure 3E**).

**Figure 3:**
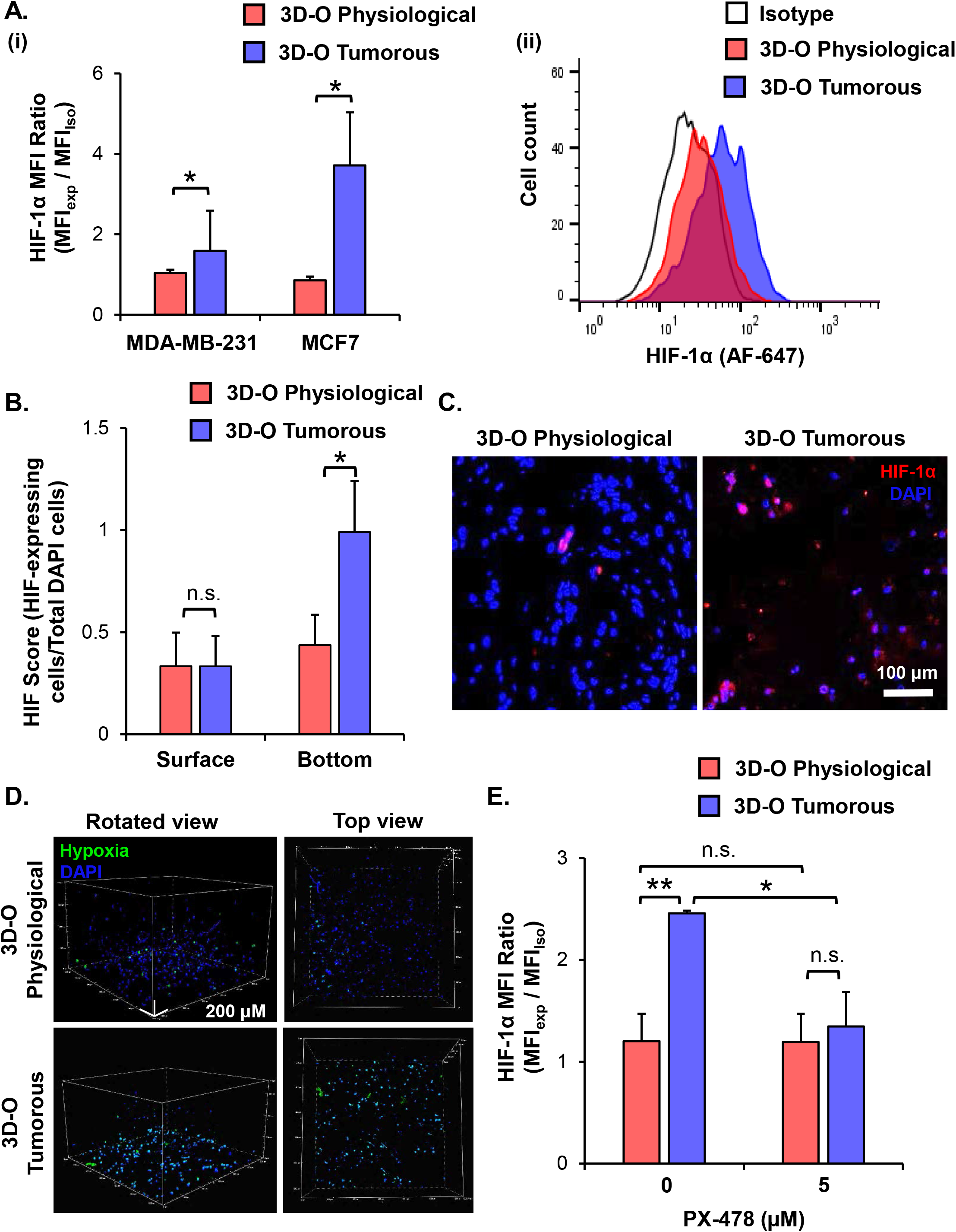
Validation of hypoxic phenotype in 3D-O matrices. A) HIF-1α expression by BCa cells grown in 3D-O physiological and 3D-O tumorous matrices after 4 days quantified as MFI ratio between AF647-anti-HIF1α and AF647 isotype control (i) and flow cytometry representative histogram (ii). B) Surface and bottom HIF-1α score indicating the percentage of BCa cells positive for HIF-1α expression after 4 days in 3D-O physiological and 3D-O tumorous matrices. C) Representative fluorescent imaging at day 4 for MDA-MB-231 cells grown within 3D-O physiological and 3D-O tumorous; DAPI: Blue; HIF-1α: Red. Merged: Pink). Scale bar= 100 μm. D) A hypoxia-sensing dye was added to both 3D-O physiological and 3D-O tumorous matrices containing MDA-MB-231 cells on day 4. After 2 hours, scaffolds were imaged using confocal microscopy to monitor hypoxia green fluorescence by resident cells. Representative fluorescent images of z-stack images at day 4 for MDA-MB-231 cells grown within 3D-O physiological and 3D-O tumorous; DAPI: Blue; Hypoxia reagent: Green). Scale bar= 200 μm. E) HIF-1α expression for MDA-MB-231 cells after treatment with PX-478 at 5 μM concentration quantified as MFI ratio between AF647-anti-HIF-1α and AF647 isotype control in 3D-O physiological and 3D-O tumorous matrices after 4 days. (**) p<0.001, (*) p<0.05, (n.s.) not significant.

### Low-oxygen within 3D-O matrices hinders BCa cell proliferation

To evaluate the impact of an oxygen-deprived environment on BCa proliferation, we analyzed BCa cell numbers in 3D-O physiological and tumorous matrices by flow cytometry. As illustrated in **Figure 4A**, for both MDA-MB-231 and MCF-7 cell lines, the rate of BCa cell proliferation was observed to be significantly hindered in 3D-O tumorous compared to 3D-O physiological model at days 4 and 7. These findings were confirmed by confocal imaging where BCa cell (DiO, green) proliferation was diminished in 3D-O tumorous compared to 3D-O physiological at days 4 and 7 (**Figure 4B**). We further corroborated a reduced BCa cell number by H&E staining and showed a remarkable reduction of Ki-67 staining, a marker of tumor cell proliferation, in 3D-O tumorous compared to 3D-O physiological (**Figure 4C**). The effect of HIF inhibitor PX-478 (5 μM) on BCa proliferation was evaluated and found that BCa cell numbers can be restored in 3D-O tumorous to the cell numbers of 3D-O physiological matrices when HIF-1α is inhibited (**Figure 4D**). These findings confirmed that BCa cell proliferation can vary depending on the oxygen content in their surrounding matrix, as mimicked by our 3D-O platforms.

**Figure 4:**
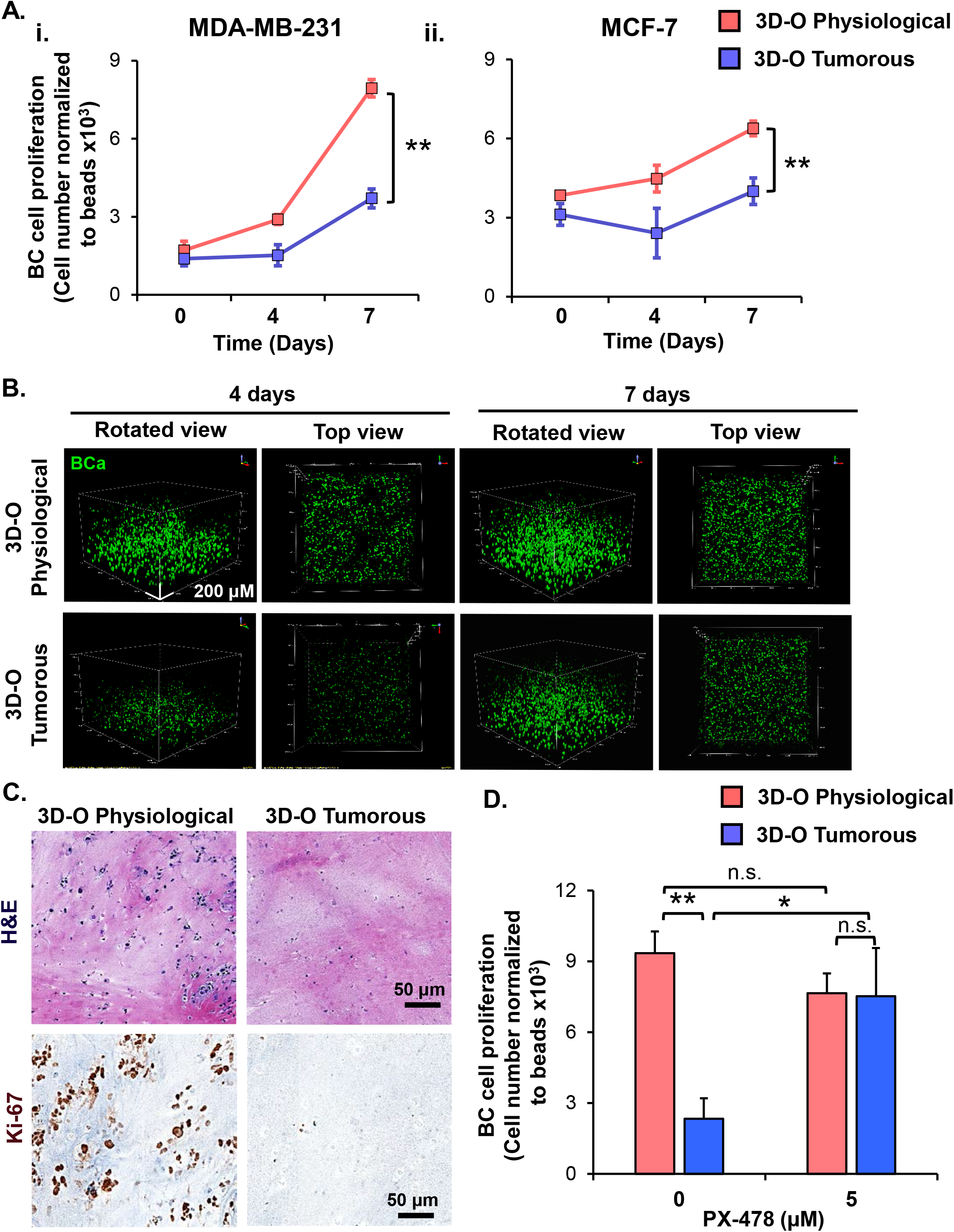
Low-oxygen within 3D-O matrices hinders BCa cell proliferation. A) Effect of oxygen deprivation on proliferation of MDA-MB-231 (i) and MCF-7 cells (ii) grown for 7 days either in 3D-O physiological or 3D-O tumorous matrices. B) Representative confocal microscopy images of MDA-MB-231 cells (green) inside 3D-O matrices at days 4 and 7 represented by z-stack images from top to bottom in rotated view and top views. Scale bar = 200 μm. C) H&E staining and IHC results representing Ki-67 expression in MDA-MB-231 cells grown for 4 days within 3D-O physiological or 3D-O tumorous matrices. Scale bar = 50 μm. D) BCa proliferation after treatment with PX-478 at 5 μM concentration quantified as cell numbers normalized to beads grown in 3D-O physiological and 3D-O tumorous matrices after 4 days, (**) p<0.001, (*) p<0.05, (n.s.) not significant.

### Low-oxygen within 3D-O matrices alters matrix environment and cellular marker expression

To characterize the role of oxygen deprivation in the surrounding matrix, we studied the expression of main fibrous ECM proteins in tissues including collagen I, collagen III, laminin and fibronectin under 3D-O tumorous and physiological conditions. Representative IF images showed a significant surge in expression for all the four studied ECMs in 3D-O tumorous as compared to the 3D-O physiological model (**Figure 5A**). Quantification of these fibrous ECM proteins indicated a significant increased expression (**Figure 5Bi**), despite a much lower BCa cell count (**Figure 5Bii**) in 3D-O tumorous compared to 3D-O physiological models. As previously observed, oxygen deprivation-driven effects can be reversed by blocking HIF-1α (PX-478, 5 μM), meaning BCa cells growing within 3D-tumorous reduced collagen I expression when HIF was inhibited (**Figure 5C**).

**Figure 5.**
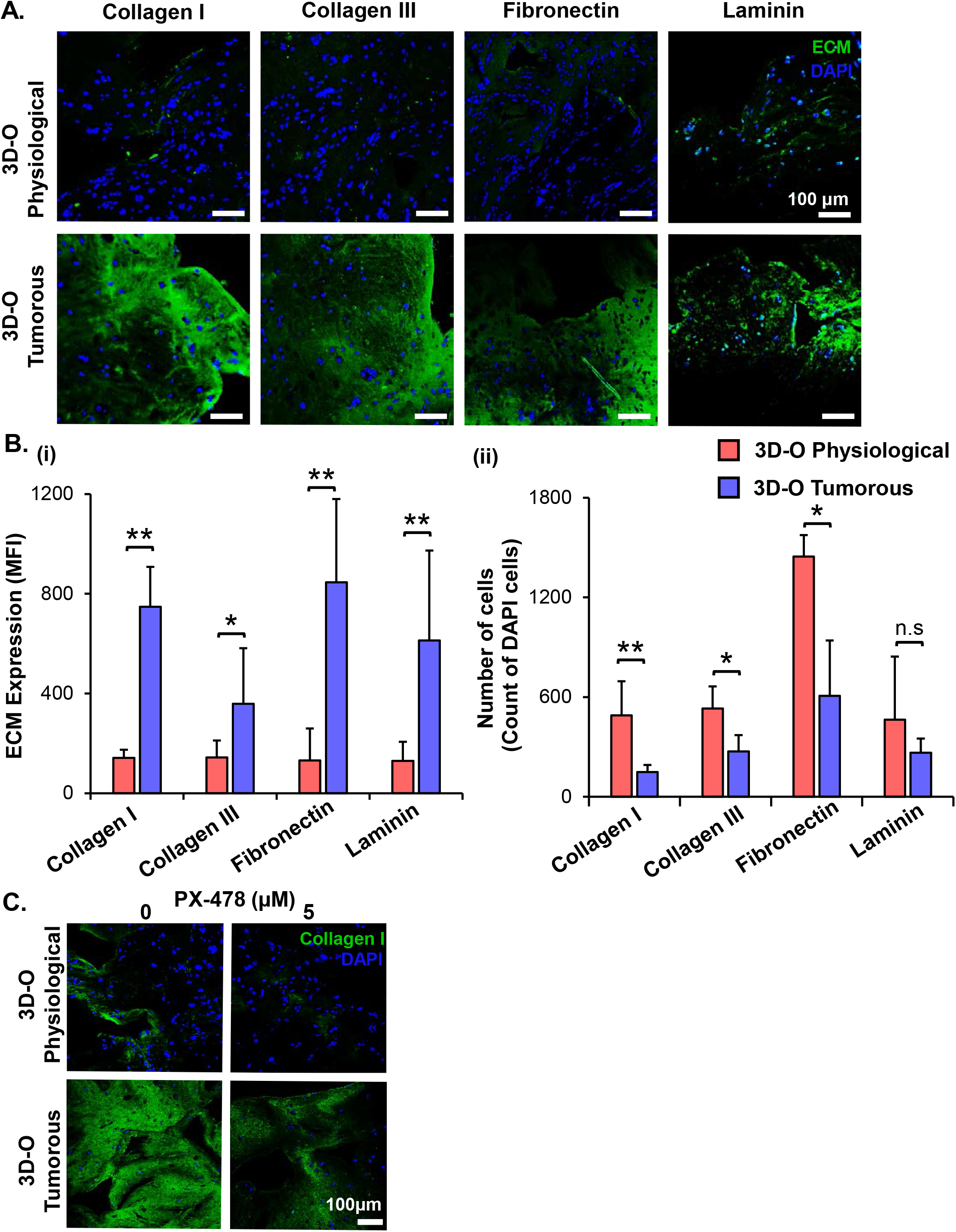
Low-oxygen within 3D-O matrices increases expression of main fibrous ECM proteins. A) Representative fluorescent images exhibiting changes in expression of collagen I, collagen III, laminin and fibronectin at day 4 for MDA-MB-231 cells grown within 3D-O physiological and 3D-O tumorous matrices; DAPI: Blue; ECMs: green). Scale bar = 100 μm. B) ECM expression by BCa cells growing within 3D-O physiological and 3D-O tumorous matrices, quantified as MFI of ECM expression (i) and number of cells (ii). C) Representative fluorescent images exhibiting changes in collagen I expression by MDA-MB-231 cells grown in 3D-O physiological and 3D-O tumorous matrices after treatment with PX-478 for 4 days at 5 μM concentration. Scale bar = 100 μm. (**) p<0.001, (*) p<0.05, (n.s.) not significant.

Additionally, we evaluated extracellular vesicle concentration under 3D-O tumorous and physiological conditions. We found that extracellular vesicle concentration under 3D-O tumorous was significantly increased compared to the 3D-O physiological model (**Suppl. Fig 1A**). Additionally, extracellular vesicle content in 3D-O models was significantly (3 and 6.7-fold) higher than conventional 2D cultures under the same incubation regimens (ambient or 1.5% O_2_), respectively (**Suppl. Fig 1B**). We further confirmed that extracellular vesicles did not show change in size (±100nm) due to oxygen deprivation or 3D culture conditions (**Suppl. Fig 1C**).

Finally, to test the effect of oxygen deprivation in BCa immune suppressive marker expression, we examined PD-L1, MUC-1 and CD73 expression on BCa cells cultured under 3D-O physiological and tumorous conditions. We found a modest increase in expression of the three immune suppressive markers in 3D-O tumorous compared to 3D-O physiological models at day 4 for both MDA-MB-231 and MCF-7 cell lines (**Suppl. Fig 2**).

### Low-oxygen within 3D-O matrices severely reduces lymphocyte infiltration

We then evaluated the effect of oxygen deprivation in lymphocyte infiltration under 3D-O physiological and tumorous conditions. We performed three independent analysis (manual gating, t-SNE and FlowSOM) to assess the effect of oxygen deprivation on lymphocyte infiltration into 3D-O models by flow cytometry. Manual gating allowed us to clearly identify infiltrated lymphocytes as CD3+ T cells (CD3-FITC+), CD4+ T cells (CD3-FITC+ CD4-PE-Cy5+), CD8+ T cytotoxic cells (CD3-FITC+ CD8-APC-Cy7+) and B cells (CD19-APC+) (**Suppl. Fig 3**).

After data processing, t-SNE and FlowSOM algorithms were used to identify the major immune cell populations in a data-driven and automated manner. t-SNE (**Figure 6A**) and FlowSOM (**Figure 6B**) assigned cells to clusters corresponding to the major immune clusters (CD3-FITC+, CD4-PE-Cy5+, CD8-APC-Cy7+, and CD19-APC+). Visualization of the high-dimensional data corresponded well to the FlowSOM algorithm (**Figure 6C**) and manual gating (**Figure 6D**).

**Figure 6:**
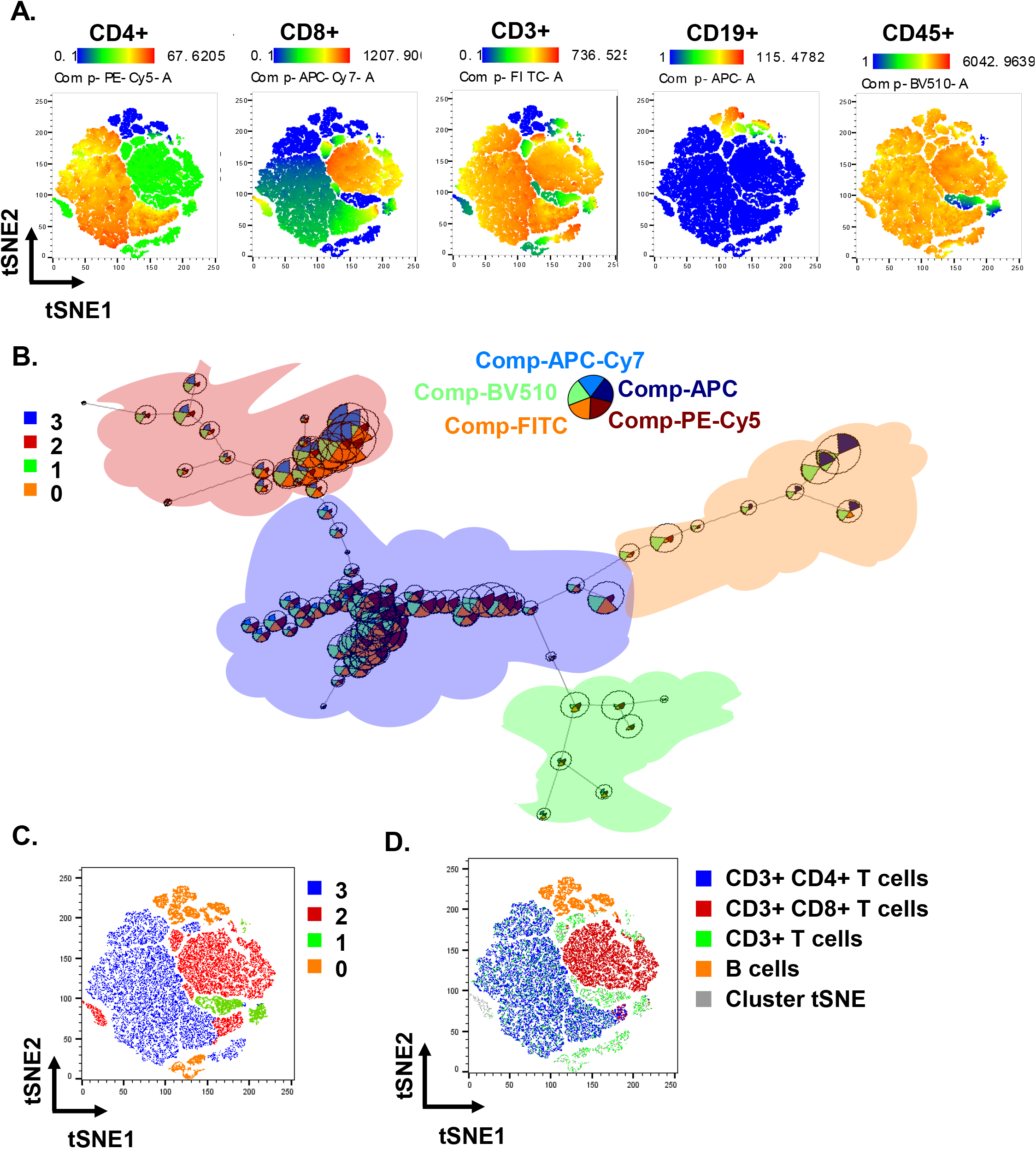
High-dimensional flow cytometry for the analysis of immune infiltration in 3D-O matrices. A) tSNE plot of pre-processed, single, live cells, down-sampled to 10.000 events and concatenated showing major immune clusters (CD3-FITC+, CD4-PE-Cy5+, CD8-APC-Cy7+, CD19-APC+ and CD45-BV510+). B) FlowSOM clustering of pre-processed, single, live cells, down-sampled to 10.000 events and concatenated revealing 4 distinct populations: 3 (BV510-high, FITC-high, PE-Cy5-high, APC-Cy7-low, APC-low), 2 (BV510-high, FITC-high, PE-Cy5-low, APC-Cy7-high, APC-low), 1 (BV510-high, FITC-high, PE-Cy5-low, APC-Cy7-low, APC-low) and 0 (BV510-high, FITC-low, PE-Cy5-low, APC-Cy7-low, APC-high). C) FlowSOM and t-SNE cluster populations are overlaid in t-SNE visualization. D) Manual gated and t-SNE cluster populations are overlaid in t-SNE visualization.

Using this approach, we identified significantly impaired infiltration of CD3+, CD4+, CD8+ and CD19+ cells inside 3D-O tumorous compared to 3D-O physiological (**Figure 7A**) by manual gating, as well as by FlowSOM (**Figure 7B**). We further confirmed impaired CD8+ infiltration by confocal imaging in 3D-O tumorous compared to 3D-O physiological (**Figure 7C**).While CD8+ infiltration in 3D-O physiological showed robust infiltration and clear co-localization of CD8+ (red) and BCa cells (green), CD8+ infiltration in 3D-O tumorous matrices was hindered.

**Figure 7:**
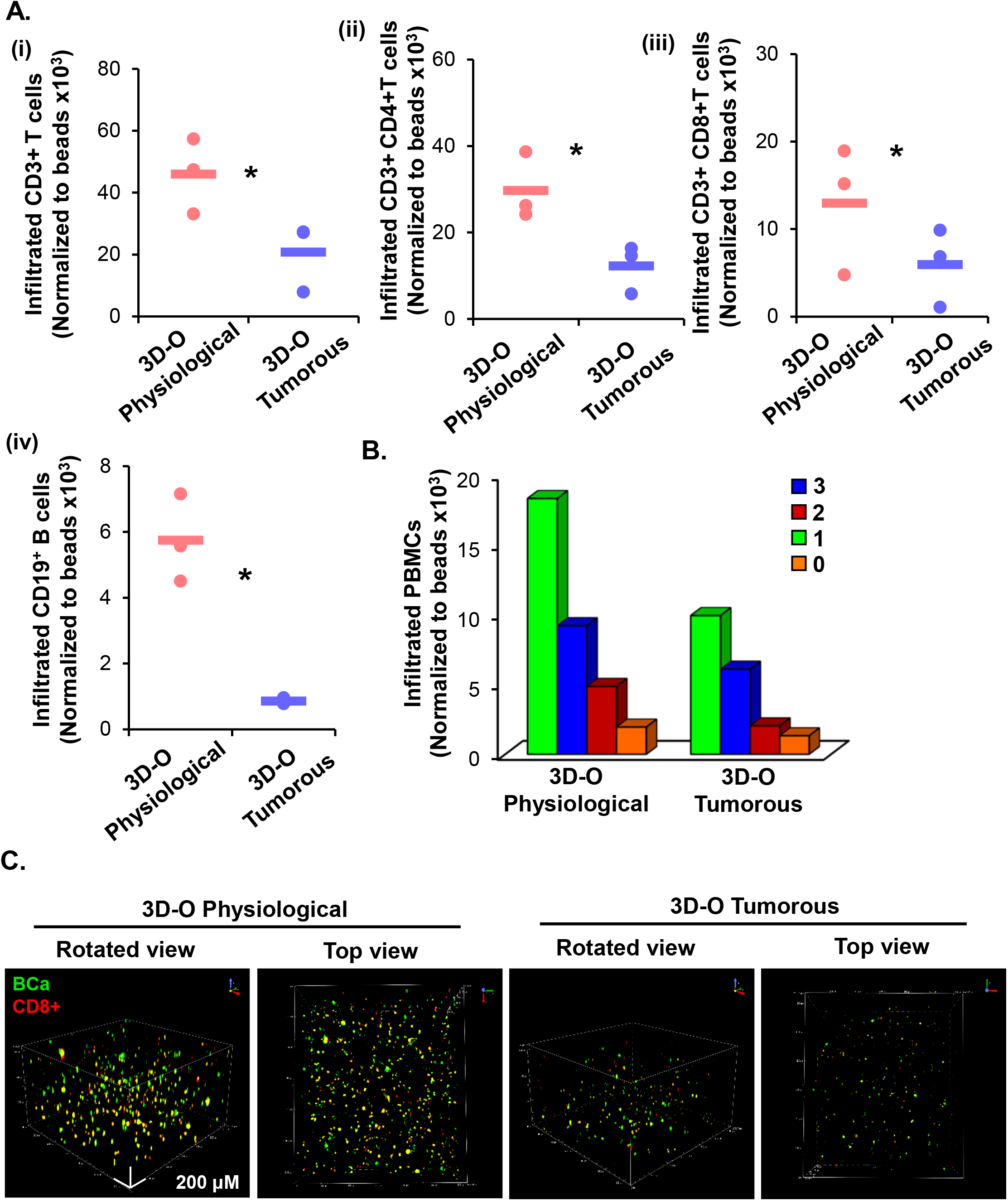
Low-oxygen within 3D-O matrices reduces lymphocyte infiltration. A) Quantification of the number of infiltrated lymphocytes into 3D-O physiological and 3D-O tumorous matrices by manual gating. Infiltration data shown represents PBMCs obtained from 3 individual healthy subjects and the average of infiltrated CD3+ T cells (i), CD3+CD4+ T cells (ii), CD3+CD8+ T cells (iii), and CD19+ B cells (data normalized to counting beads ×10^3^), (*) p<0.05 B) Quantification of the cell numbers across infiltrated populations into 3D-O matrices using FlowSOM. C) Representative confocal microscopy images of rotated and top views of BCa (green) cells and CD8+ (red) cells in 3D-O physiological and 3D-O tumorous matrices at day 7, co-localization is seen in yellow. Scale bar = 200 μm.

Moreover, the effect of HIF inhibitor PX-478 (5 μM) on CD8 infiltration was evaluated and found that CD8+ infiltration can be restored in 3D-O tumorous to the cell infiltration numbers of the 3D-O physiological model when HIF-1α is inhibited (**Figure 8A**). These findings were corroborated by IHC staining for CD8 (**Figure 8B**). Additionally, we found that treatment with an investigational anti-PD-L1 monoclonal antibody (Durvalumab, 5μM) did also reverse CD8 infiltration in 3D-O tumorous to the cell infiltration numbers of the 3D-O physiological matrices (**Figure 8C**). This finding correlates with our previous result showing that the expression of PD-L1 was enhanced in 3D-O tumorous matrices (**Suppl. Fig 2A**).

**Figure 8:**
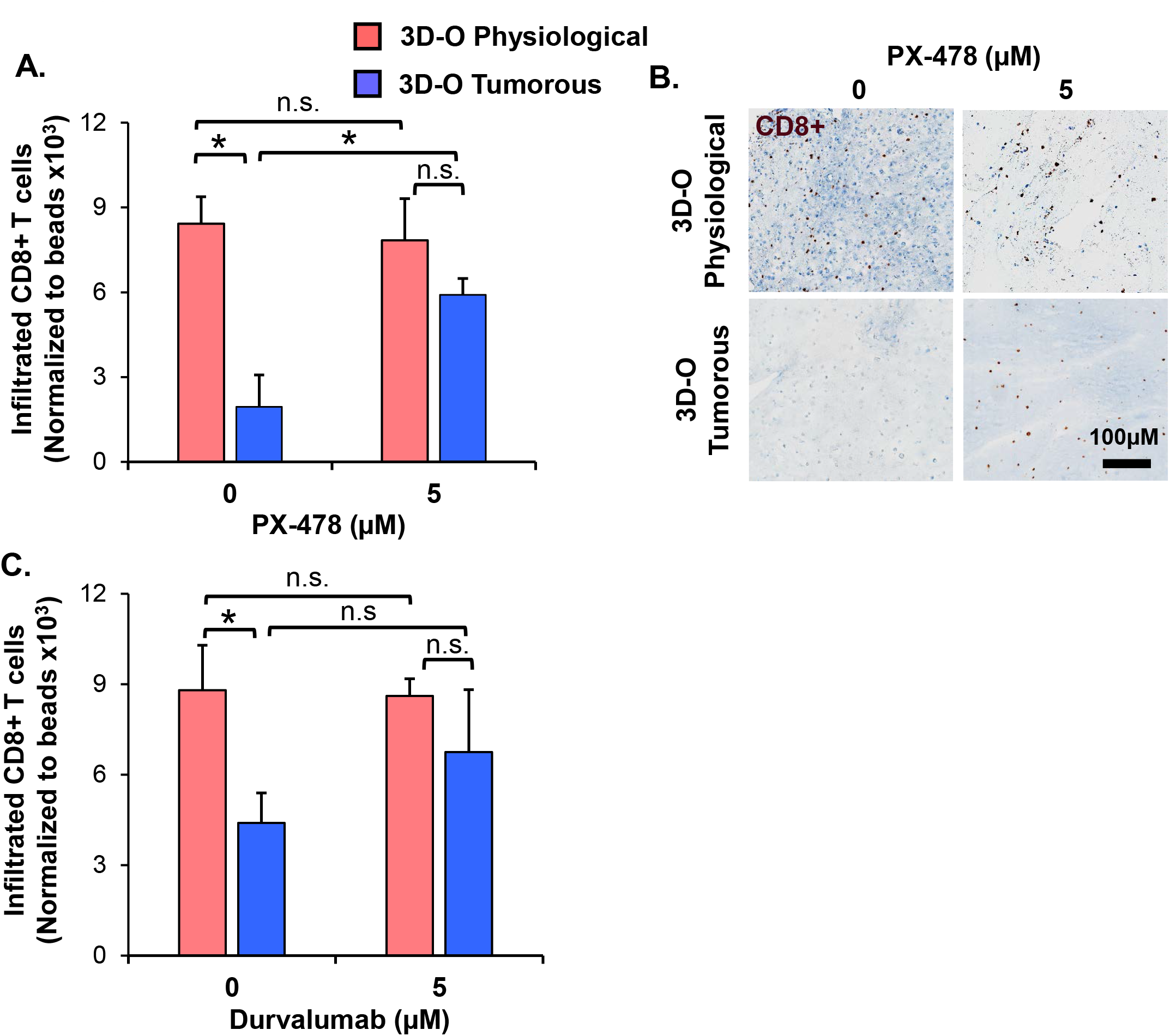
Sensitization of BCa cells to cytotoxic CD8+ T cells within 3D-O matrices. A) CD8+ infiltration into 3D-O physiological and 3D-O tumorous matrices on day 7 after treatment with PX-478 at 5 μM concentration for the first 4 days. B) Representative IHC results representing CD8^+^ infiltration into 3D-physiologial and 3D-tumorous scaffolds on day 7 after treatment with PX-478, Scale bar = 100 μm. C) CD8^+^ infiltration into 3D-O physiological and 3D-O tumorous matrices on day 7 after treatment with Durvalumab at 5 μM concentration for first 4 days. (*) p<0.05, (n.s.) not significant.

## DISCUSSION

An adaptive, key hallmark of cancer is the tumor’s capacity to accommodate to changeable states of oxygen deprivation. As oxygen content regulation is essential to maintaining cell homeostasis, small deviations in oxygen level in the TME can result in major changes in tumor cell functionality (34, 58–60). Physiological oxygen levels in solid tumors are very heterogeneous ranging from 0.5 – 4% oxygen saturation compared to 4 – 14% in healthy tissues (59, 61–63). In addition, circulating and immune cells are exposed to a full dynamic range of blood physiological O_2_, in which the pO_2_ varies from 13.5% in alveoli and 9.5% in arterial blood to 6.5% at the venous end of circulation (64–68). Therefore, studies employing models incubated at atmospheric oxygen (21% O_2_) culture conditions are hyperoxic with respect to the tissue of origin (65, 69). Oxygen fluctuation occurs at irregular intervals in cancer with sporadic reoxygenation periods, but undoubtedly oxygen depletion in tumors directly affects clinical responses to therapy by influencing tumor growth and is associated with a more aggressive tumor phenotype (70–72). Mounting evidence from cancer research demonstrates that oxygen deprivation significantly promotes stark differences in cancer behaviors including tumor progression, therapy resistance and immunosuppression (73–76). To better understand the role of the tissue microenvironment on variations of cancer development it is necessary to account for oxygen levels originating from the tumor and from microenvironment interactions. Therefore, the use of models that can recreate physiologically applicable oxygen conditions might be more useful for research purposes.

The challenges involved with translation of bench therapies to clinical outcomes perhaps originate from limitation of suitable models to adequately mimic the native oxygen deprived TME in its entirety (77–79). Recreating a relevant oxygen profile in current cancer models might be essential for determining tumor-immune evasive mechanisms. Very few models have been recruited to decipher the interactions involved in oxygen deprivation and immune evasion inside the tumor. Immune cell infiltration into 3D matrices has been observed to vary significantly based on the oxygen content, when cancer cells and highly efficient chimeric antigen receptor T (CAR-T) cells were incorporated *in-vitro* in a 3-D micro pattern within a photo-crosslinked hydrogel and coupled to a microfluidic hypoxia device (Ando et al., 2019). In this regard, we developed a gelatinous-like and porous 3D model, where 3D-O physiological with 14–10.9 kPa pO_2_ recreates physiological oxygen levels of healthy tissue and blood physiological levels that circulating and immune cells are exposed to (29, 62) and 3D-O tumorous with 9 – 2 kPa pO_2_ (29, 30, 62) recreates pathophysiological oxygen levels of tumor tissue (**Figures 1 and 2**).

The 3D-O model is developed from naturally occurring patient-derived resources minimizing the introduction of foreign elements that can alter the natural pathophysiology of the TME. The oxygen content within these 3D-O matrices not only modifies according to the surrounding culture conditions, but it recapitulates several tumor hypoxia hallmarks known to be critical targets for designing novel therapies. There are several advantages of developing a platform with plasma as the framework. Being enriched with growth factors and proteins that contribute to the regenerative process, a plasma derived microenvironment potentiates cell proliferation, migration, and differentiation mimicking *in vivo* conditions, hence enhancing the clinical utility of the model (80–83). Of note, the use of human plasma as a matrix not only supports cell structure, but also serves as a reservoir for nutrients, growth factors, cytokines, extracellular vesicles, and signaling molecules (84–86) that help with the development of more relevant human cancer models. Utilizing plasma is very cost effective and helps conserve accessibility and safety by using the patient’s own intrinsic constituents. In the last few years, many 3D models have employed plasma-based biomaterials, with both structural and functional purposes (41, 83, 86–88).

Our studies demonstrated that the 3D-O model while supporting BCa cell growth, simulated physiological oxygen levels and generated intratumoral hypoxic hallmarks such as increased HIF expression (**Figure 3**). Upregulated HIF expression is a well-known parameter associated with hypoxia and is known to affect several downstream biochemical processes (89). Our results confirm that oxygen deprivation within 3D-O can efficiently reiterate HIF-driven regulation in the resident BCa cells. It is well known that within the tumor hypoxic niche, a high level of HIF-driven adaptations are observed in terms of ECM modeling (90), expression of immune surface markers PD-L1 (91), MUC-1 (92), CD73 (93) and hypoxia-induced extracellular vesicles (94). Low oxygen-induced changes within the 3D-O supported many known tumor hypoxia characteristics such as reduced BCa cell proliferation and alteration of the TME such as increased ECM protein expression, increased extracellular vesicle secretion and enhanced immune surface marker expression of BCa cells (**Figures 4, 5 and Suppl. Fig 1 and 2**). In addition, 3D-O showed that there was a higher amount of secreted extracellular vesicles generated as a result of the three-dimensional environment when compared to 2D and these changes were further enhanced in response to oxygen deprivation. Extracellular vesicle yield in 3D cultures has already been shown to be higher than in 2D models (95), as well as that hypoxic conditions can lead cells to express increased vesicle yield (96). The increased expression for each of the studied ECMs in the presence of lower oxygen could either be accounted for by an absolute increase in new ECM generation or by an enhanced enzymatic functional cascade resulting in enhanced alignment of the ECMs already present in the matrix environment. But overall, the different ECM levels by BCa cells growing within our 3D-O model reflect the importance of these changes with respect to reported literature about how ECMs can promote and maintain BCa tumor health and cancer progression (97).

BCa is characterized by a highly inflammatory microenvironment supported by infiltrating immune cells, cytokines, and growth factors (98, 99). Moreover, the importance of boosting TILs in disease management has been cited as a critical therapeutic strategy (100). Among TILs, CD8^+^ T cells have been shown to migrate and infiltrate tumors and mediate antitumoral responses (101). Significant crosstalk exists between hypoxia and antitumor immune functions, where tumor hypoxia contributes to attenuated antitumor responses (102). There is evidence that oxygen deprivation influences many immune cells, including T cells and B cells (103). HIF-1α expression in cancer cells has been linked to reduce ability of CD8+ T cells to recognize malignancy and lower the ability of T cells to proliferate (104). The ability to mimic tumor-mediated immune escape mechanisms driven by oxygen deprivation could be functionally fundamental for selecting the best strategies to combine therapies that have the maximum potential to fully eradicate cancers (105). We further demonstrated that low oxygen-induced changes mimicked tumor-immune interactions leading to immune evasion mechanisms. CD8+ T cells infiltration was significantly impaired under pathophysiological oxygen levels and further confirmed that HIF and PD-L1 inhibition, sensitize BCa cells to cytotoxic CD8+ T cells (**Figures 7 and 8**). Many previous studies have addressed managing the hypoxic tumor using HIF and PD-L1 inhibitors which resulted in promising outcomes (91, 106). It is well established that boosting the CD8+ cell infiltration into the tumor niche can reverse immune evasion (107–110). Results from our experiments using PX-478 and Durvalumab on 3D-O matrices confirm that reversing lymphocyte functional impairment is possible by either targeting HIF-1α or targeting inhibitory checkpoint pathways like PD-L1/PD-1.

The 3D-O platform can serve as a model that bridges the gap between tumor hypoxia, immune evasion and recreating TME features as close as possible to *in vivo* conditions. The 3D-O model will allow us to study novel targeted therapies to try to overcome hypoxia-driven immune evasion. The clinical value of assessing oxygen deprivation will reveal how hypoxia-modification therapy could benefit patients with hypoxic tumors (111–113). As hypoxia within the tumor is the cause of several therapeutic discrepancies, a hypoxia specific approach would perhaps be practical to study tumor-immune interactions. Further studies will characterize the functionality of infiltrated immune populations within the oxygen-deprived environment. Additional studies where patient-isolated cancer and accessory cells from a tumor biopsy are incorporated into 3D-O matrices to assess the depth-driven oxygen effects in the presence of immunomodulatory drugs and TME components would also be needed. We believe that the results from these studies would greatly improve the understanding of how immune cells adapt to low-oxygen microenvironments.

In conclusion, the 3D-O model exhibited physio- and pathophysiological oxygen levels that allowed us to further characterize low oxygen-induced changes in BCa including impaired cell proliferation, altered TME, and hindered immune infiltration. The 3D-O model could serve as a promising platform for the evaluation of immunological events and as a drug-screening platform tool to overcome hypoxia-driven immune evasion.

## ACKNOWLEDGEMENTS

We want to thank Erin Harmon and Daniel S. Engebretson from the Biomedical Engineering Department at the University of South Dakota for their help with SEM. We also want to thank Kelly Graber from the Histology and Imaging Core at Sanford Research for her training and help with the Imaging. This project was supported by an Institutional Development Award (IDeA) from the National Institute of General Medical Sciences of the National Institutes of Health under grant number 5P20GM103548 and the 2018 LUSH Prize (Young Investigator Award). This project used Sanford Research Histology and Imaging Core, and Flow Cytometry Core Facilities that are supported in part by a COBRE grant from the National Institutes of Health (P20 GM103548-06).

## DATA AVAILABILITY STATEMENT

The raw/processed data that support the findings of this study are available from the corresponding author, PP, upon reasonable request.

## AUTHORSHIP

Somshuvra Bhattacharya: methodology, investigation, validation, formal analysis, visualization, writing - original draft and writing -review & editing. Kristin Calar and Mark Petrasko: investigation, validation and writing - review & editing. Claire Evans: resources and writing - review & editing. Pilar de la Puente: conceptualization, methodology, visualization, writing - original draft and writing - review & editing, supervision, project administration and funding acquisition.

