## Supplemental data for "Bioengineering a novel 3D in-vitro model to recreate physiological oxygen levels and tumor-immune interactions"

1 Cancer Biology and Immunotherapies Group, Sanford Research, Sioux Falls, SD, USA.
2 Histology and Imaging Core, Sanford Research, Sioux Falls, SD, USA.
3 Sanford PROMISE, Sanford Research, Sioux Falls, SD, USA.
4 Department of Surgery, University of South Dakota Sanford School of Medicine, Sioux Falls, SD, USA.
5 Department of Chemistry and Biochemistry, South Dakota State University, Brookings, SD, USA.
6 Flow Cytometry Core, Sanford Research, Sioux Falls, SD, USA.

Short title: Bioengineering oxygen levels and tumor-immune interactions

* Corresponding Author:

Pilar de la Puente, Ph.D., M.Sc.

Assistant Scientist

Cancer Biology and Immunotherapies Group

Sanford Research

2301 E 60th Street N

Sioux Falls, SD 57104

#### FIGURE LEGENDS

**Supplementary Figure 1: Low-oxygen within 3D-O matrices reduces extracellular vesicle secretion.** A) Concentration of extracellular vesicles generated per MDA-MB-231 cell (ng/mL) grown within 3D-O physiological and 3D-O tumorous matrices for 4 days. B) Fold change concentration of extracellular vesicles generated per cell (ng/mL) grown for 4 days either in 3D-O culture or matching classic 2D culture and incubated either in ambient air or 1.5% $O_2$. C) Representative plot for extracellular vesicle size (nm) versus concentration (particles/ml). (**) $p<0.001$, (*) $p<0.05$.

**Supplementary Figure 2: Low-oxygen within 3D-O matrices moderately enhances immune surface marker expression.** A) Immune surface marker expression by BCa cells grown in 3D-O physiological and 3D-O tumorous matrices after 4 days quantified as MFI ratio for PD-L1 (i), CD73 (ii), and MUC-1 (iii). B) Flow cytometry representative histograms for PD-L1 (i), CD73 (ii), and MUC-1 (iii).

**Supplementary Figure 3: Manual flow cytometry gating strategy for the analysis of immune infiltration in 3D-O matrices.** Representative 2D dot plots demonstrating gating strategy for defining major lymphocyte populations infiltrated in 3D-O matrices. Data acquisition was performed with a fixed number of counting beads as constant With this strategy, the first gate excludes all cellular populations from beads, followed by gating singlets. Then, lymphocyte populations were gated based on forward and side scatter (FSC and SSC) and the live cells (Live/Dead viability marker) were identified. Following this, CD3+ T cells (CD3-FITC+), CD4+ T cells (CD3-FITC+ CD4-PE-Cy5+), CD8+ T cytotoxic cells (CD3-FITC+ CD8-APC-Cy7+) and B cells (CD19-APC+) were gated from the live lymphocyte population. The CD8 by CD4 gate helps distinguish true $CD8^+$ or $CD4^+$ cells and excludes the double negative or positive auto fluorescent cells. Gating on the fluorescence minus one (FMO) controls were used to set gates for FITC (CD3), PE-Cy5 (CD4), APC-Cy7 (CD8), APC (CD19) and BV510 (CD45). FMOs for each test color were used as controls.

**Suppl. Fig 1.**

**A.**

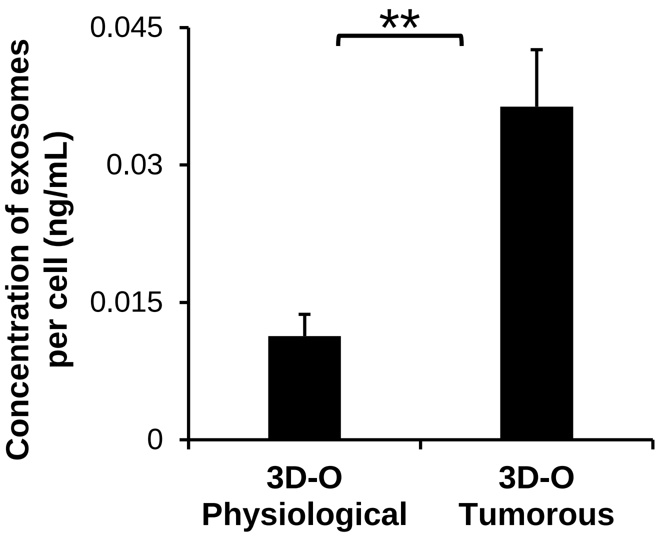


**B.**

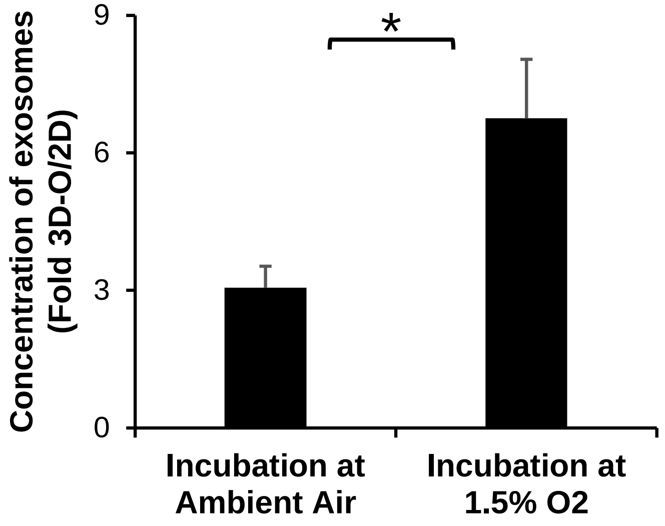


**C.**

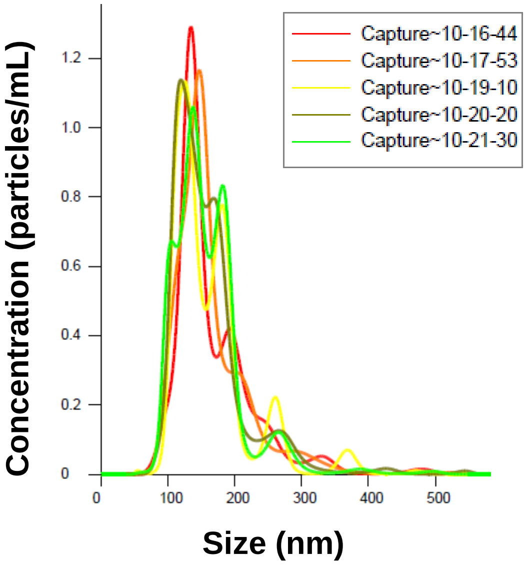

Suppl. Fig 2.

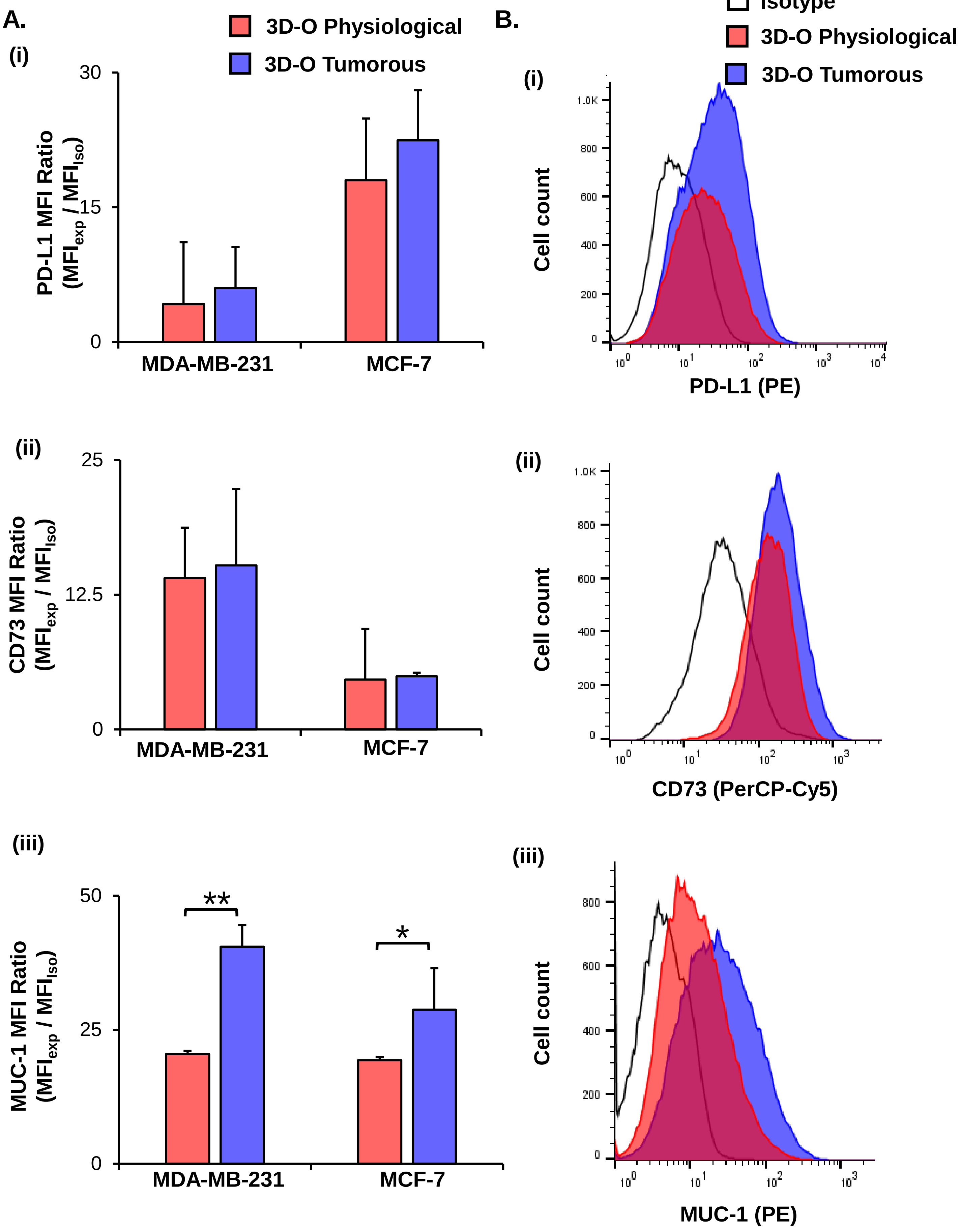
A.
B.
3D-O Physiological
3D-O Tumorous
Isotype
(i)
PD-L1 MFI Ratio
(MFIexp / MFIIso)
30
15
0
MDA-MB-231
MCF-7
Cell count
PD-L1 (PE)
(ii)
CD73 MFI Ratio
(MFIexp / MFIIso)
25
12.5
CD73 (PerCP-Cy5)
(iii)
MUC-1 MFI Ratio
(MFIexp / MFIIso)
50
25
**
*
MUC-1 (PE)

### Suppl. Fig 3.

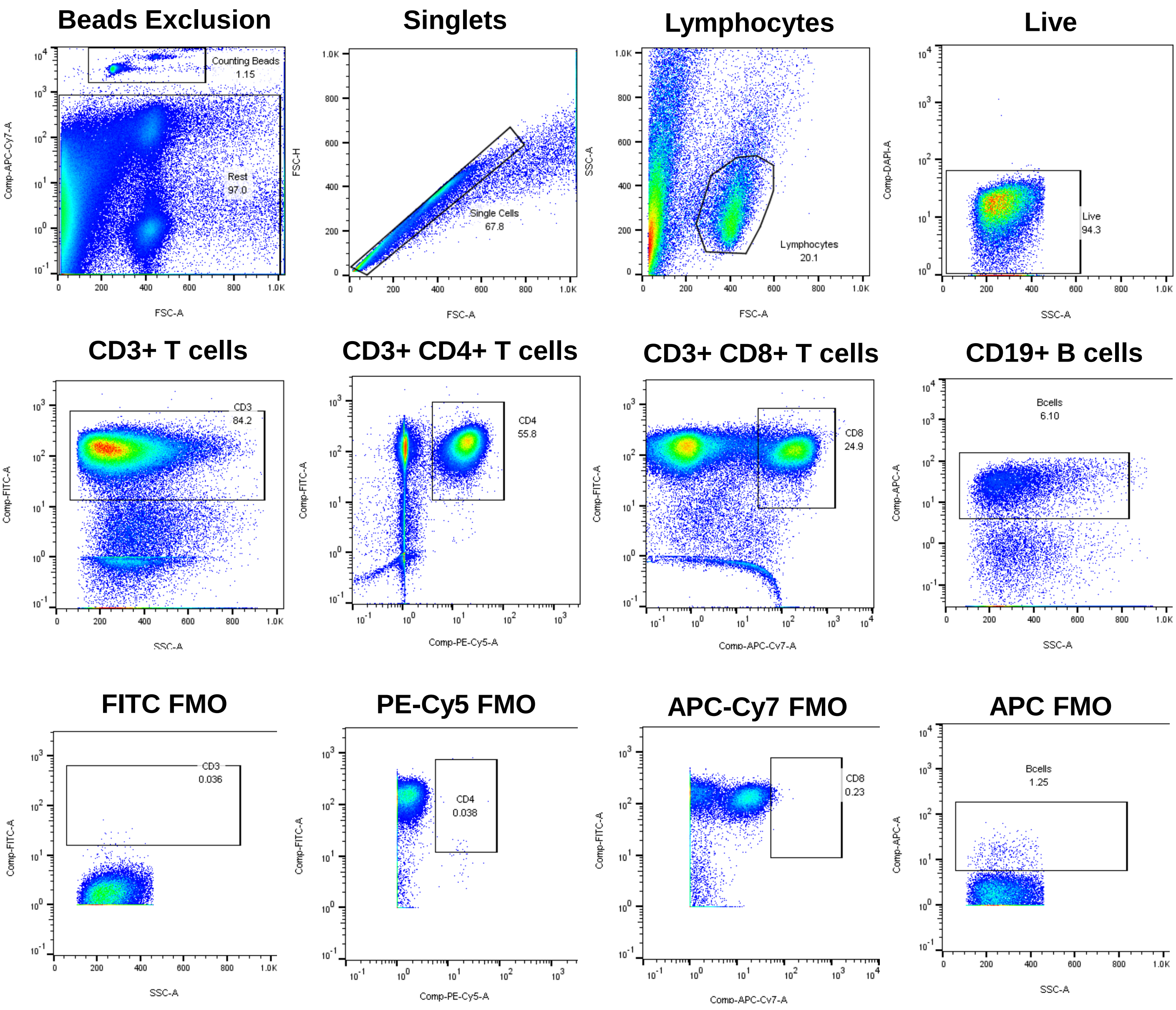
Beads Exclusion
Counting Beads
1.15
Rest
97.0
Comp-APC-Cy7-A
FSC-A
Singlets
Single Cells
67.8
FSC-H
FSC-A
Lymphocytes
Lymphocytes
20.1
SSC-A
FSC-A
Live
Live
94.3
Comp-DAPI-A
SSC-A
CD3+ T cells
CD3
84.2
Comp-FITC-A
SSC-A
CD3+ CD4+ T cells
CD4
55.8
Comp-FITC-A
Comp-PE-Cy5-A
CD3+ CD8+ T cells
CD8
24.9
Comp-FITC-A
Comp-APC-Cy7-A
CD19+ B cells
Bcells
6.10
Comp-APC-A
SSC-A
FITC FMO
CD3
0.036
Comp-FITC-A
SSC-A
PE-Cy5 FMO
CD4
0.038
Comp-FITC-A
Comp-PE-Cy5-A
APC-Cy7 FMO
CD8
0.23
Comp-FITC-A
Comp-APC-Cy7-A
APC FMO
Bcells
1.25
Comp-APC-A
SSC-A
